# Sexually dimorphic differences in angiogenesis markers predict brain aging trajectories

**DOI:** 10.1101/2023.07.16.549192

**Authors:** A Torres-Espin, Hannah Rabadaugh, S Fitzsimons, D Harvey, A Chou, C Lindberg, KB Casaletto, L Goldberger, AM Staffaroni, P Maillard, BL Miller, C DeCarli, JD Hinman, AR Ferguson, JH Kramer, FM Elahi

**Author notes:** **Corresponding Author:** Fanny M. Elahi, MD PhD Annenberg 20-50, Box 1137 One Gustave L. Levy Place New York, NY 10029. These authors contributed equally (A Torres-Espin, Hannah Rabadaugh).

## Abstract

Aberrant angiogenesis could contribute to cognitive impairment, representing a therapeutic target for preventing dementia. However, most angiogenesis studies focus on model organisms. To test the relevance of angiogenesis to human cognitive aging, we evaluated associations of circulating blood markers of angiogenesis with brain aging trajectories in two deeply phenotyped human cohorts (n=435, age 74±9) with longitudinal cognitive assessments, biospecimens, structural brain imaging, and clinical data. Machine learning and traditional statistics revealed sex dimorphic associations of plasma angiogenic growth factors with brain aging outcomes. Specifically, angiogenesis is associated with higher executive function and less brain atrophy in younger women (not men), a directionality of association that reverses around age 75. Higher levels of basic fibroblast growth factor, known for pleiotropic effects on multiple cell types, predicted favorable cognitive trajectories. This work demonstrates the relevance of angiogenesis to brain aging with important therapeutic implications for vascular cognitive impairment and dementia.

## INTRODUCTION

The aging of vasculature represents an integral component of the organ-based age-associated vulnerability to disease(*1*). With aging, numerous pathologies result in abnormal blood vessels across calibers of vessels from capillaries to large vessels(*2*). The pathologies present in small blood vessels are among the most insidious, and yet prevalent and detrimental consequences of aging for systemic homeostasis, including immune system regulation, and brain function(*3, 4*). Of these age-related microvasculopathies, cerebral small vessel disease (cSVD), has the strongest association with cognitive impairment(*5*). Prevalent and cSVD-associated cognitive changes include slowed processing speed and decline in executive function(*6*). The constellation of cognitive impairment and cerebrovascular disease is referred to as vascular cognitive impairment (VCI)(*7*), implying suspected causality, with numerous aspects of cSVD being potent drivers of brain dysfunction and degeneration(*4*). Therefore, molecular mechanisms underpinning cSVD represent an active area of research and therapeutic development. In chronic cSVD related VCI, several multi-cellular pathologies have emerged, such as capillary rarefaction(*8, 9*), dysfunctional BBB(*10, 11*), impaired neurovascular coupling(*12*), brain hypoperfusion, neuroinflammation, and myelin degeneration and impaired repair(*5*). Notwithstanding, most discoveries have been made in model systems. The molecules involved in these diverse pathologies and the pathways linking the many pathological aspects of cSVD with cognition in humans remain in large part poorly understood. Currently, the diagnosis and quantification of severity of cSVD pathologies related to cognitive impairment remain challenging in humans(*13*), and in contrast to Alzheimer’s disease, there is an unmet need for molecular biomarkers of VCI.

One emerging molecular pathway important for the establishment and maintenance of healthy vasculature, including healthy capillaries, is angiogenesis(*14*). This is also a molecular pathway that could potentially link dysfunctions in development(*15*) to selective vulnerability in brain degeneration. Angiogenesis, the highly regulated process of new blood vessel formation, is fundamental to health across organ systems and an important molecular pathway for healthy aging(*1*), with brain dysfunction and cognitive impairment representing brain phenotypes in the setting of systemic dysregulations. Several studies found a decline in the density of capillaries with age, known as capillary rarefaction, accentuated in those with neurodegenerative disease or in carriers of risk genes(*8, 16*). A study has shown that countering the age-associated impairment in angiogenesis can increase health-span in mice(*1*), with an effect across all organs investigated. This age-associated decline in angiogenesis appears to be multifactorial, with contributions from changes in alternative splicing and localization of membrane bound receptor, FLT1 (Fms Related Receptor Tyrosine Kinase 1, also known as VEGFR), to soluble FLT1 (sFLT1) which acts as a VEGFA trap, interfering with its angiogenic function(*17*). There may be additional phenotypic changes of endothelial cells contributing to impaired microvascular function and lack of appropriate angiogenic response in aging, such as loss of endothelial cellular identity and function with age leading to the development of phenotypes such as increased non-specific vascular permeability along with concomitant decline in selective transport(*18*).

Based on the critical role of angiogenesis for all organs, it is plausible to hypothesize that in the brain, dysregulations in angiogenesis and molecular signaling that maintain microvascular structures and function beyond development may play important roles in brain aging and cognitive impairment. Still, there is an important gap between clear mechanistic studies in model systems, which demonstrate the importance of angiogenesis to brain aging pathologies, and translational studies of brain aging in humans. The combination of molecular markers of angiogenesis with age-pertinent structural and functional (cognitive) outcomes is an important first step to bridging this gap and understanding whether angiogenesis could be a target for brain aging therapeutics, worth pursuing in clinical trials. We tested for associations of circulating blood markers of angiogenesis with brain aging trajectories in two deeply phenotyped aging cohorts with longitudinal cognition, biospecimen, structural brain imaging and clinical data using unsupervised machine learning linear mixed effects models. The results surprisingly reveal sexually dimorphic and age-dependent associations between markers of angiogenesis and brain aging outcomes.

## RESULTS

### The trajectory of plasma markers of angiogenesis is associated with aging

#### Temporal covariance profiles of angiogenesis

We quantified our primary markers of interest, Phosphatidylinositol Glycan Anchor Biosynthesis Class F (PlGF), FLT1, and Fibroblast Growth Factor 2 or Basic Fibroblast Growth Factor (bFGF), along with other proteins using a commercially available, technically reliable hypersensitive multi-plex approach that included a total of seven markers of angiogenesis (PlGF, FLT1, Tie2, VEGFA, VEGFC, VEGFD and bFGF) with the Meso Scale Discovery (MSD) V-Plex electrochemiluminescence assay (Meso Scale Diagnostics, Rockville, Maryland). The antibodies used to detect FLT1 with this assay are not isoform specific and could detect both the secreted form (sFLT1) as well as the solubilized membrane-bound protein. We used an unsupervised machine learning workflow through missing data analysis and robust non-linear principal component analysis (NL-PCA) to uncover multivariate patterns of association between these markers, leverage data richness, and limit multiple comparisons. With this approach, we identified temporal covariance profiles of angiogenesis proteins (markers) in the panel of seven markers in the form of principal components (PCs) (**Fig. 1**). Two significant and robust patterns of marker associations (PCs) emerged by a permutation-based test of the variance explained by each component (Torres-Espin et al., 2021; Linting et al., 2011) (**Fig. 1e**). Results show that PC1 explains 24.9% of the variance in the dataset and is robustly loaded by PlGF, FLT1, VEGFA and VEGFC with approximately the same contributions in the same direction (**Fig. 1f**). PC2 explains 18.8% of the variance and is loaded by bFGF, VEGFA and VEGFC in one direction, and loaded by PlGF, FLT1, and Tie2 in the inverse direction. Notably, bFGF was the highest positive-loading marker in PC2 (**Fig. 1g**). The most striking difference between these PCs is an implied inverse relationship of our primary markers of interest. More specifically, PlGF and FLT1 both load positively onto PC1 whereas FLT1 and PlGF load inversely to bFGF onto PC2 (**Supplementary Fig. 1**). Furthermore, while both PlGF and FLT1 load highly onto PC1, bFGF clearly emerges as a strong driver of PC2. This may suggest that bFGF levels capture a biological process that differs from those captured by PlGF and FLT1. Finally, we studied the robustness of these patterns with a resampling method of bootstrapping NL-PCA 500 times, resulting in highly similar PC1 and PC2 to the ones described above (average coefficient of congruence of 0.997 and 0.998 for PC1 and PC2 respectively, **Supplementary table 1**). These results suggest that our derived measures are stable and indicative of reproducibility and generalizability of uncovered patterns.

**Figure 1.**
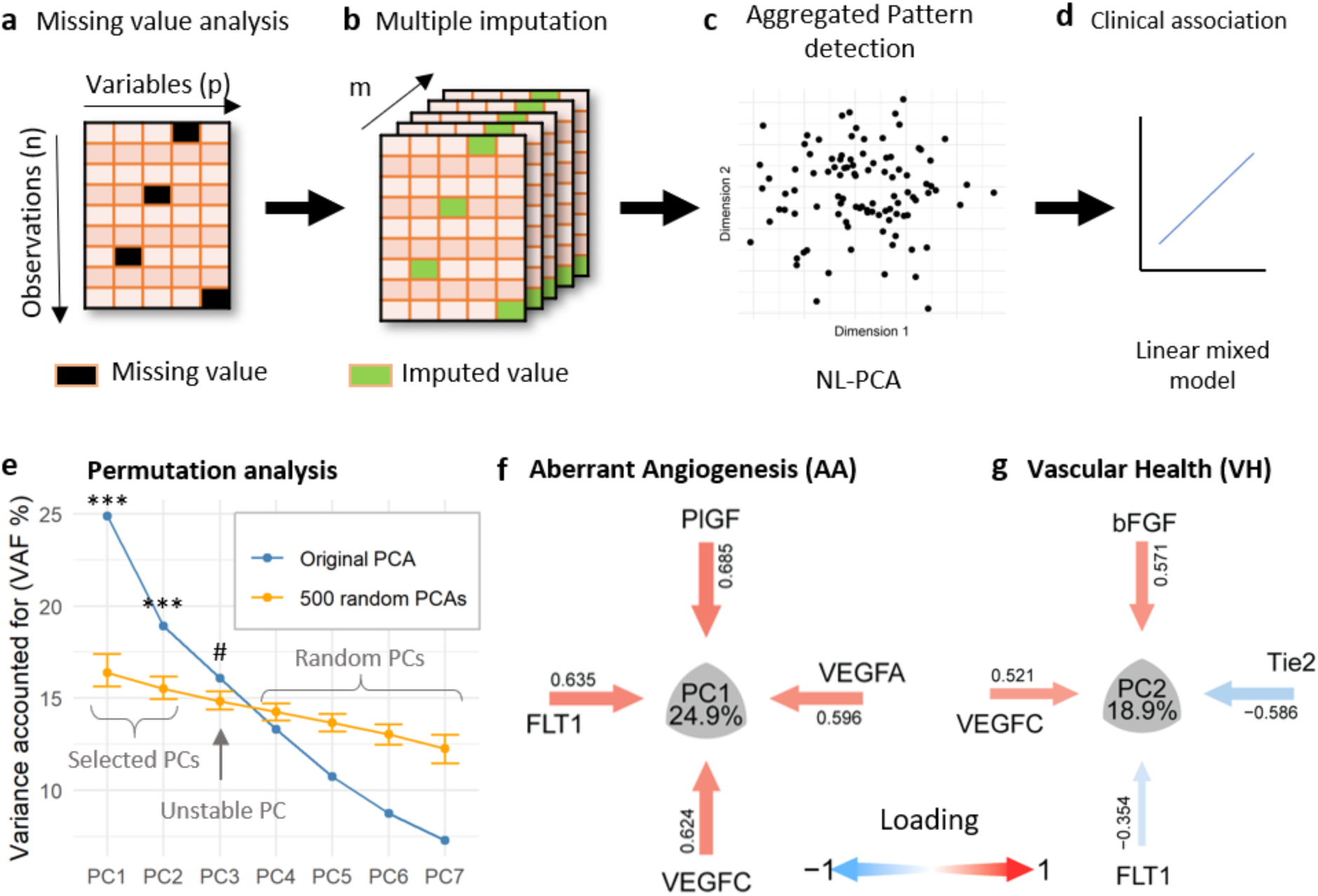
Unsupervised angiogenesis pattern detection workflow. Missing values analysis was performed to determine data missing patterns across all observations (n) and variables (p) (**a**). Missing values were then imputed using multiple imputations by chained equations (MICE) producing m number of imputed datasets, in this study m = 20 (**b**). The resulting imputed datasets were aggregated using the median for each data point and a non-linear principal component analysis (NL-PCA) was performed to extract patterns of association or principal components (PC) between variables in a low dimensional space (**c**). The resulting PC scores were used for further analysis of clinical association by linear mixed models (**d**). The number of relevant PCs were selected using a permutation-based method. Only PCs from the orinigal NL-PCA (**a** to **c**) with information exceeding that of randomly generated PCs, loaded on by more than one variable and stable to missigness were retained (**e**). PC1, renamed Aberrant Angiogenesis (AA), explained 24.9% of the variance (**f**), and PC2, renamed Vascular Health (VH), explained 18.9% of the variance (**g**).

#### Angiogenesis and Aging Trajectories: Associations with Demographics

With the overarching hypothesis that plasma concentrations of molecular drivers of angiogenesis captured by PC1 and PC2 may mark pathological aging trajectories, we began by testing the associations of PC1 and PC2 with chronological age, modelling in the effects of sex and *APOE* genotype as pertinent biological variables to brain aging. Specifically, we hypothesized that PC1 and PC2 would be associated with chronological age as this measure is the best available surrogate for biological age. Given the scarcity of information regarding the role of angiogenesis in brain aging and neurodegenerative disease, we did not know whether *APOE4* carrier status would have an effect nor what the effect of sex would be, but, if these markers act on AD-pertinent pathological pathways, an interaction with *APOE4* and sex could be conceivable, as carriers, especially women carriers, have higher risk of symptomatic brain degeneration with AD neuropathology(*19*).

We found that PC1 increases with age in women more than men and that PC2 decreases with age in both sexes (**Fig. 2**). Based on this initial result, we formulated the following specific hypotheses that we tested. We hypothesized that increasing levels of PC1 reflect pathological aspects of angiogenesis in aging, and that decreasing levels of PC2 correlate with decline in healthy brain physiology with age. As such PC1 levels would be associated with declining measures of brain health (cognition and brain volumes) and the reverse would be expected for PC2. Thus, we renamed PC1 Aberrant Angiogenesis (AA) and PC2 Vascular Health (VH) and will refer to them as such throughout this manuscript. The directionality of change with age of the individual markers of interest were found to be congruent with PCs. Both PlGF and FLT1 demonstrate an increase with age, while bFGF demonstrates a decrease with age (**Fig. 2**). We did not find evidence of sex differences when considering these markers individually (**Supplementary tables 4-6**).

**Figure 2.**
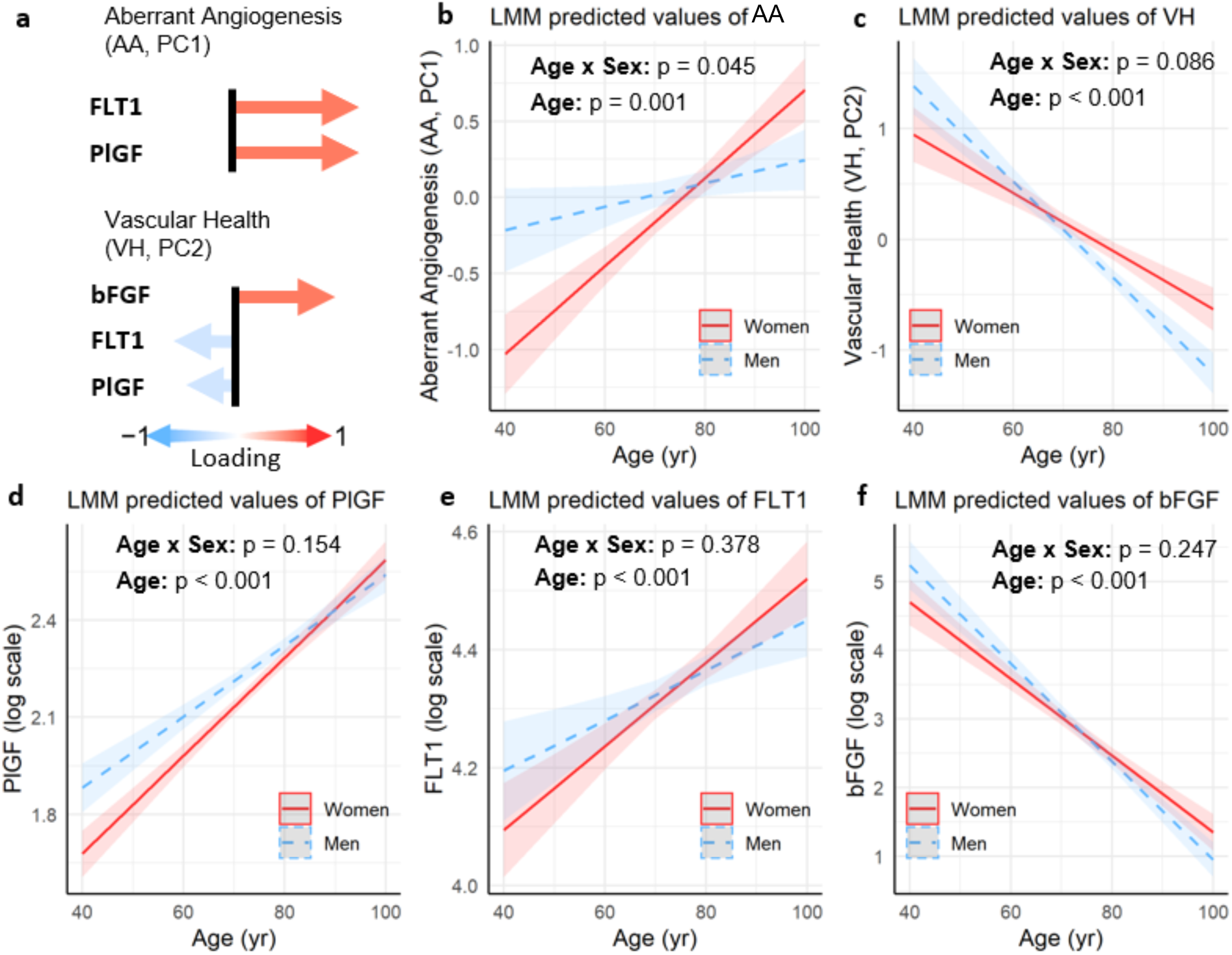
Demographic associations with angiogenic patterns. The contribution of markers of interest (PlGF, FLT1, bFGF) with each of the selected PCs (Aberrant Angiogenesis, PC1; Vascular Health, PC2) is shown in (**a)**. Arrows pointing to the right (red) represent positive loadings and arrows pointing to the left (blue) represent negative loadings. The relationship of each PC scores and each marker levels to age and sex has been studied using linear mixed models (LMM). The graphs show the predicted value (bold line) and standard error (shadow ribbon) of PC1(AA) (**b**), PC2 (VH) (**c**), PlGF (**d**), FLT1 (**e**) and bFGF (**f**). For each model, the p value for the Age x Sex interaction term and for the Age term is provided.

Next, considering the higher risk for Alzheimer’s disease neuropathological changes in women carriers of *APOE4*(*20*), we wondered whether the associations of these markers with age differed by *APOE* genotype and sex. We found that age-associated changes in AA and VH were *APOE* genotype-independent (**supplementary tables 2, 3**).

#### Cerebrovascular disease and risk factors are associated with plasma levels of angiogenic growth factors

Since angiogenesis protein markers were quantified from plasma, the levels could reflect to pathophysiology and disease of any vascular bed, including but not exclusive to brain and heart. We built statistical models to test the association of PCs and select plasma markers of interest with vascular risk factors. Systemic vascular risk factors included hypertension, hyperlipidemia, and diabetes. In our dataset, clinical diagnoses pertinent to brain and heart vascular disease, such as stroke and myocardial infarction, were clearly captured. We again took an unbiased approach for dimension reduction, applying NL-PCA to derive and retain three vascular principal components (vPC) (Supplementary **Fig. 2**). We find that the first vascular principal component (vPC1) mainly captures variance among vascular risk factors, while the second and third vascular principal components (vPC2 and vPC3, respectively) capture variance of cardiovascular and cerebrovascular disease burdens, respectively. These diagnoses are elicited through a history obtained by a clinician during the participant’s research visit. PC loadings indicate that while cardiovascular disease loaded on vPC2, vascular risk factors had minimal contribution to this PC. This finding may be due to an indication bias such that risk factors detected early would have been treated and therefore reducing burden of future disease. Furthermore, since history-based vascular risk and disease-associated diagnoses may be impacted by participant recall bias, we also assessed associations of plasma angiogenic markers with systolic blood pressure measured at the in-person visit for participants, which is exempt from recall bias. Of note, blood pressure is also considered to be one of the most important risk factors for VCID(*21*).

In the cohorts for this study, vascular risk factors captured by vPC1 increases across age in women, while cerebrovascular disease burden, captured by vPC3, increases across age in men (Supplementary **Fig. 2**). These sex differences may be reflecting recruitment bias or diagnostic bias and provide additional information regarding vascular risk in our cohort. No age associated change is noted for vPC2, the cardiovascular PC. With respect to individual markers, levels of bFGF inversely correlate with vascular risk factors (vPC1) in younger individuals in the cohort and positively associated with vPC1 in older individuals, with a transition in direction of the association noted around age 75 (Supplementary **Fig. 3**, **Table 2**). While a directional turning point was only noted for vPC1, levels of bFGF were inversely associated with the cardiovascular disease burden (vPC2) regardless of sex and age. These results indicate that higher levels of bFGF, whether a reflection of health state or potential compensatory increase to counteract age-associated disease, may predict better brain aging trajectories. We found FLT1 levels to have a positive association with vPC2 and vPC3 in men (Supplementary **Fig. 3, Table 2)**. Finally, no significant associations were noted between vPCs and PlGF.

**Table 1.** Study Participant Demographics. Mean, median for demographics and cognition and percentage of missing data by sex. CDR=Clinical Dementia Rating.

|  | Men (N=228) | Women (N=207) | Overall (N=435) |
| --- | --- | --- | --- |
| **# visits** |  |  |  |
| Mean (SD) | 6.01 (4.31) | 5.78 (4.21) | 5.90 (4.26) |
| Median [Min, Max] | 5.00 [1.00, 20.0] | 5.00 [1.00, 19.0] | 5.00 [1.00, 20.0] |
| **Mean Age** |  |  |  |
| Mean (SD) | 75.2 (8.71) | 73.4 (8.79) | 74.3 (8.78) |
| Median [Min, Max] | 75.9 [46.8, 92.5] | 74.0 [46.6, 92.3] | 75.0 [46.6, 92.5] |
| **Age at first visit** |  |  |  |
| Mean (SD) | 72.2 (8.59) | 70.6 (8.57) | 71.4 (8.61) |
| Median [Min, Max] | 72.5 [46.8, 89.9] | 70.9 [42.6, 88.4] | 71.7 [42.6, 89.9] |
| **Age at last visit** |  |  |  |
| Mean (SD) | 78.0 (9.50) | 76.2 (9.79) | 77.1 (9.67) |
| Median [Min, Max] | 79.0 [46.8, 100] | 76.9 [47.5, 98.2] | 77.9 [46.8, 100] |
| **APOE E4 status** |  |  |  |
| Negative | 116 (50.9%) | 127 (61.4%) | 243 (55.9%) |
| Positive | 78 (34.2%) | 53 (25.6%) | 131 (30.1%) |
| Missing | 34 (14.9%) | 27 (13.0%) | 61 (14.0%) |
| **Mean Executive function** |  |  |  |
| Mean (SD) | -0.198 (0.723) | -0.137 (0.710) | -0.169 (0.717) |
| Median [Min, Max] | -0.255 [-2.56, 1.84] | -0.114 [-2.38, 1.60] | -0.174 [-2.56, 1.84] |
| Missing | 15 (6.6%) | 9 (4.3%) | 24 (5.5%) |
| **Executive function first available visit** |  |  |  |
| Mean (SD) | -0.159 (0.713) | -0.127 (0.699) | -0.144 (0.706) |
| Median [Min, Max] | -0.193 [-2.56, 2.04] | -0.106 [-2.38, 1.75] | -0.144 [-2.56, 2.04] |
| Missing | 15 (6.6%) | 10 (4.8%) | 25 (5.7%) |
| **Executive function last available visit** |  |  |  |
| Mean (SD) | -0.271 (0.802) | -0.200 (0.763) | -0.237 (0.783) |
| Median [Min, Max] | -0.328 [-2.72, 1.81] | -0.122 [-2.58, 1.40] | -0.232 [-2.72, 1.81] |
| Missing | 15 (6.6%) | 10 (4.8%) | 25 (5.7%) |
| **Mean CDR sum of boxes** |  |  |  |
| Mean (SD) | 1.69 (2.19) | 1.25 (1.99) | 1.48 (2.10) |
| Median [Min, Max] | 0.577 [0, 12.3] | 0.200 [0, 13.2] | 0.400 [0, 13.2] |
| Missing | 0 (0%) | 2 (1.0%) | 2 (0.5%) |
| **CDR sum of boxes first available visit** |  |  |  |
| Mean (SD) | 1.27 (2.12) | 0.995 (1.77) | 1.14 (1.96) |
| Median [Min, Max] | 0.500 [0, 17.0] | 0 [0, 13.0] | 0 [0, 17.0] |
| Missing | 15 (6.6%) | 10 (4.8%) | 25 (5.7%) |
| **CDR sum of boxes last available visit** |  |  |  |
| Mean (SD) | 2.23 (3.25) | 1.42 (2.33) | 1.84 (2.87) |
| Median [Min, Max] | 0.500 [0, 17.0] | 0 [0, 14.0] | 0.500 [0, 17.0] |
| Missing | 15 (6.6%) | 10 (4.8%) | 25 (5.7%) |

**Table 2.**
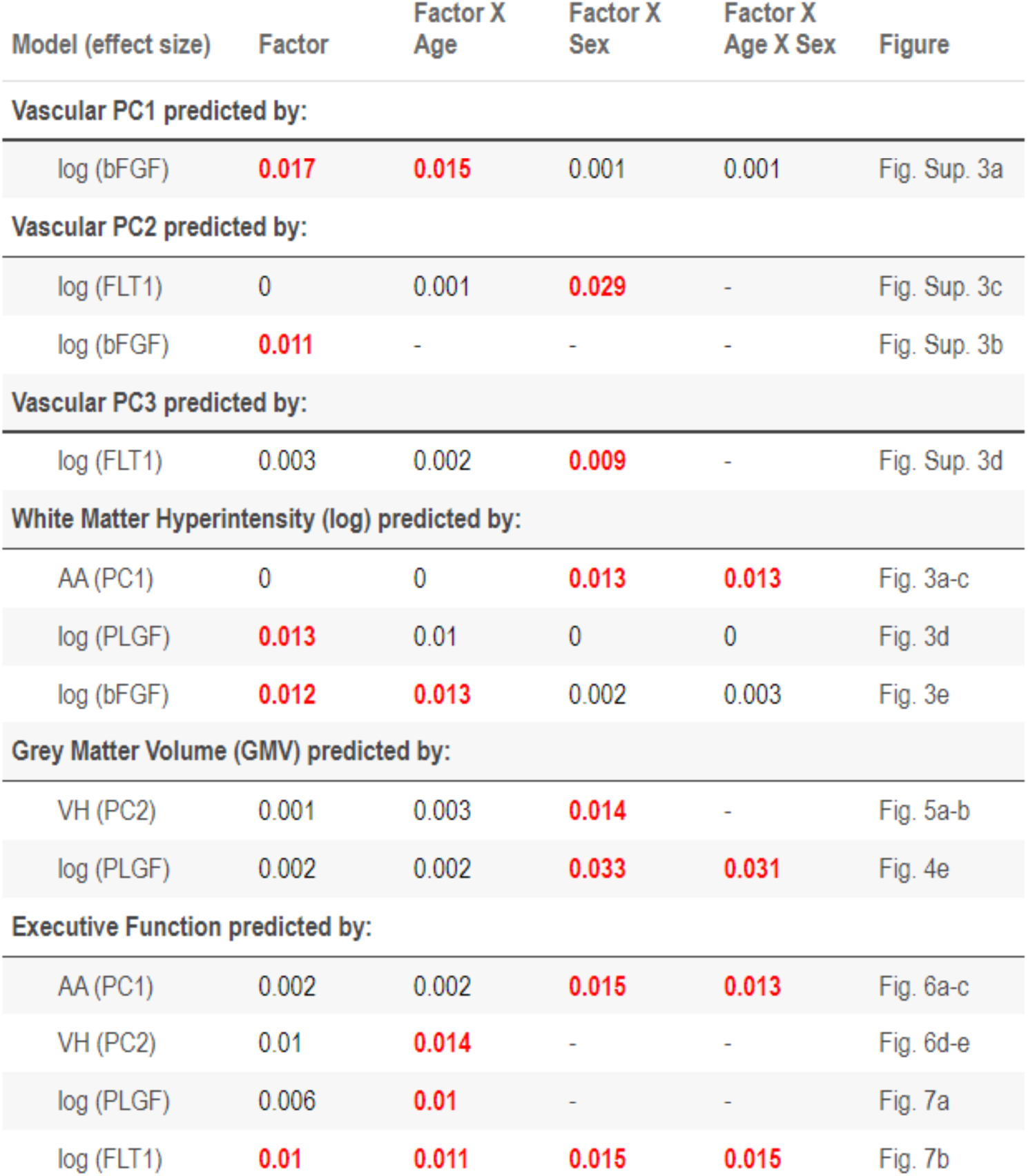
Summary statistics of interpreted models with age and gender associations. Effect size (partial eta square) is shown for each association, with significant (p < 0.05) effects marked in red. “-“ indicates that the specific term as not tested. X indicates interaction. Factor makes reference to the specific predictor denoted in the first column

### Plasma markers of angiogenesis are associated with brain health trajectories

As discussed earlier, we found that plasma measures of AA, as well as PlGF and FLT1, increase with chronological age while VH and bFGF decrease with chronological age. Considering these results, we evaluated the “pathological” nature of elevation in AA, PlGF and FLT1 levels by testing associations with measures of pathological brain aging such as white and grey matter atrophy (surrogate for neurodegeneration), white matter injury (white matter hyperintensity, WMH), and decline in cognition, with a focus on executive function as the most pertinent cognitive domain for VCID.

#### Neuroimaging markers of vascular brain injury

We found that AA is positively associated with burden of white matter hyperintensity (WMH). The association of AA with WMH is sex and age-dependent, with higher levels of AA associated with higher WMH in older women, while an inverse association is noted in younger women, congruent with findings discussed above. Interestingly, in men the direction of association was inversed, such that higher levels of AA are associated with lower WMH in older men and higher WMH in younger men (**Fig. 3a-c**).

**Figure 3.**
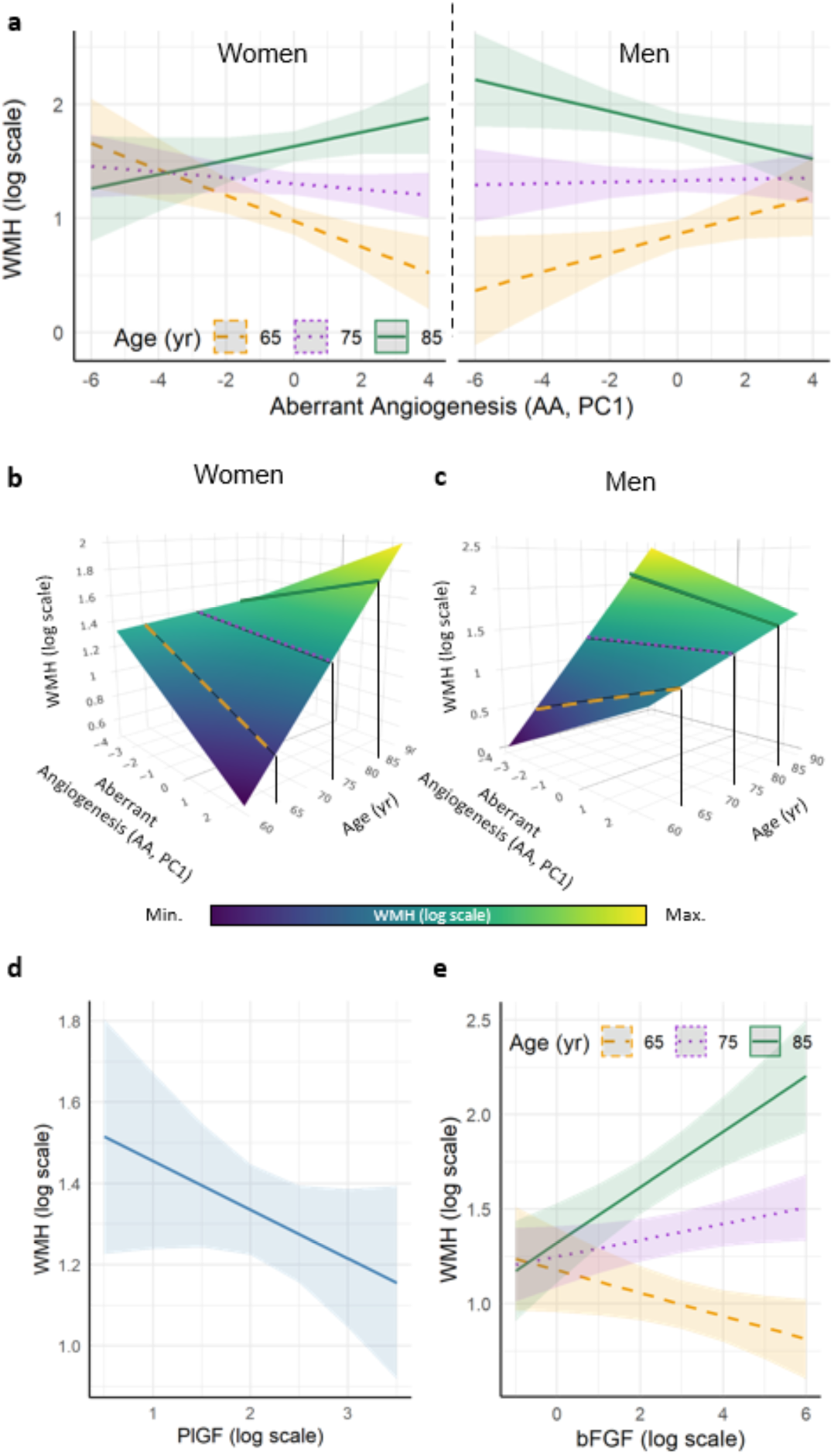
Association of angiogenic principal components with neuroimaging marker of white matter injury. Aberrant angiogenesis associates with markers of vascular brain injury. Linear mixed models (LMM) were used to study the association of AA with white matter hyperintensity (WMH) when considering the effect of age and sex. We found an association of PC1 to WMH values dependent on age and sex (PC1 x age x Sex p = 0.03). To visualize all the effects described by the model, we represent the predicted values of WMH (**a**) for women and men separately. An illustrative representation of the continuous dependent effect of age on PC1 association with WMH, the prediction for three ages (65, 75 and 85 years) are shown in the line graphs (**a**). Considering all potential predictions of the model given age creates a prediction surface in a 3D space (**b** and **c**). The three ages are shown over the surface for reference. The interactive plot for **b** and **c** can be found on the supplementary material (supplementary interactive figures). PlGF had a significant effect on explaining WMH (p = 0.035) (**d**). The model predicting WMH showed a statistically significant interaction between bFGF levels and Age (p = 0.028) (**e**). The graphs show the predicted value (bold line or surface) and standard error (shadow ribbon).

Although VH is not associated with WMH, bFGF demonstrates an age-dependent association with WMH, such that higher bFGF is associated with higher WMH in older ages, starting at 71-72 (**Fig. 3e**). This relationship does not appear to be dependent on sex or *APOE* genotype (**Supplementary table 19**).

#### Plasma angiogenesis markers are associated with neurodegeneration in aging

The neuroimaging measure of WMH can broadly capture numerous cellular pathologies, including glial activation, BBB permeability, interstitial edema, as well as atrophy and degeneration of neuronal axons and microvasculature. Notwithstanding, the most proximal neuroimaging marker of cognitive impairment remains neuronal degeneration captured by grey matter volume (GMV) loss or GM atrophy. Therefore, GMV is a good, classical surrogate marker for neurodegeneration(*22*). When investigating associations of angiogenic markers with GMV (**Fig. 4 and 5**, **Table 3**), our findings are conceptually congruent with findings reported above. We find an age-dependent relationship between AA and GMV, such that higher levels of AA are associated with higher GMV in younger women and transition to be an inverse association with GMV in older women, irrespective of *APOE* status. In older men, a positive association between AA and GMV is noted in *APOE4* carriers. Again, we note that higher levels of AA may reflect and predict higher measures of brain health in older men (**Fig. 4a**, **Table 3**), while higher levels appear to be associated with lower measures of brain health in older women (**Fig. 4a**, **Table 3**)

**Figure 4.**
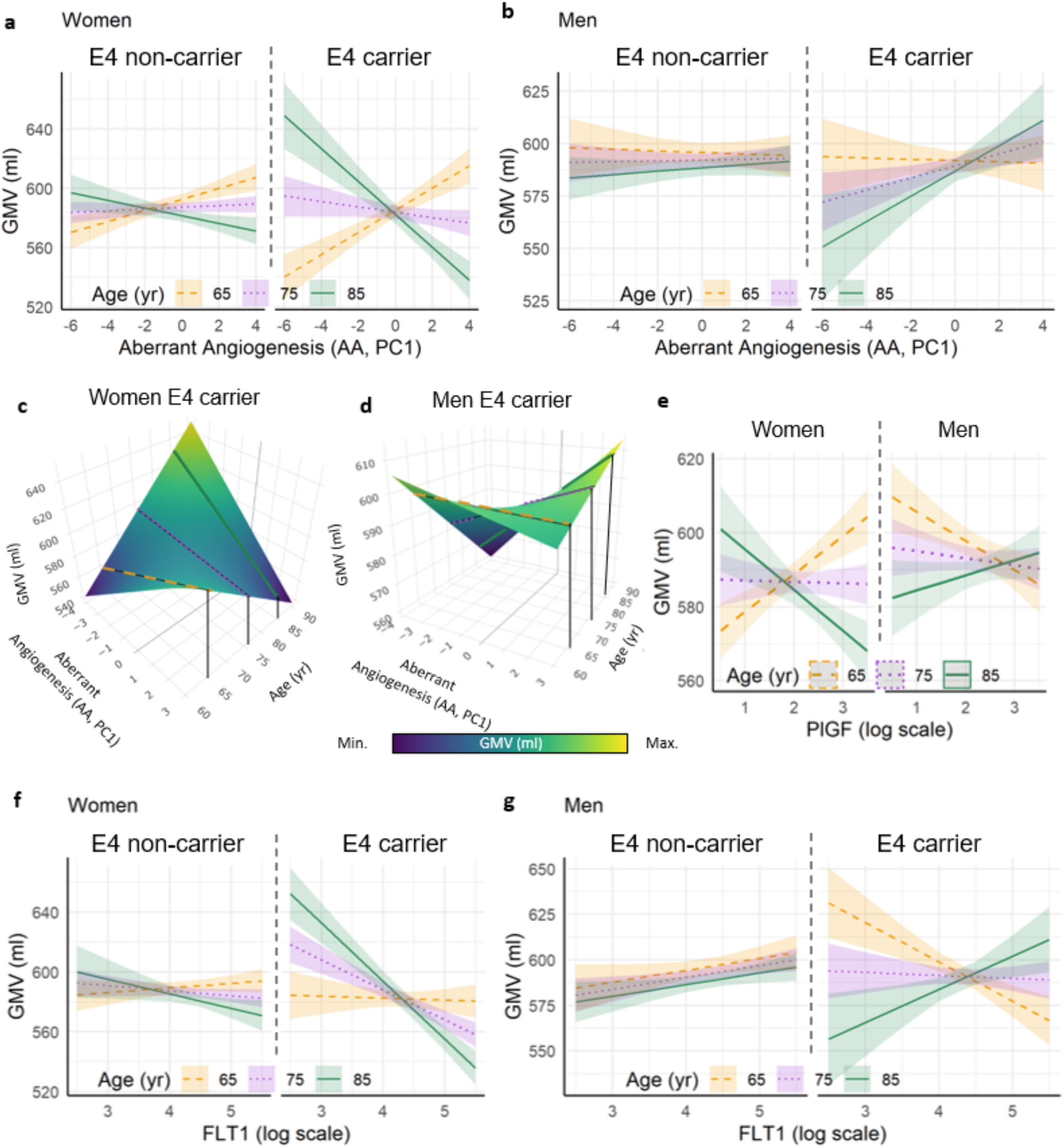
Association of Aberrant Angiogenesis to grey matter volume. The LMM analysis that models the association of AA with GMV shows a significant interaction between AA, age, sex and APOE genotype (p = 0.033). The continuous interaction of AA and age for E4 non-carriers and carriers Women and for Men is shown in **a** and **b** respectively. The prediction for three ages (65, 75 and 85 years) are shown in the line graphs. Considering all potential predictions of the model given age creates a prediction surface in a 3D space (**b** for E4 carrier women; **c** for E4 carrier men). The interactive plot for **a** and **b** can be found on the supplementary material (supplementary interactive figures). The graphs show the predicted value (bold line or surface) and standard error (shadow ribbon). See also Table 3.

**Figure 5.**
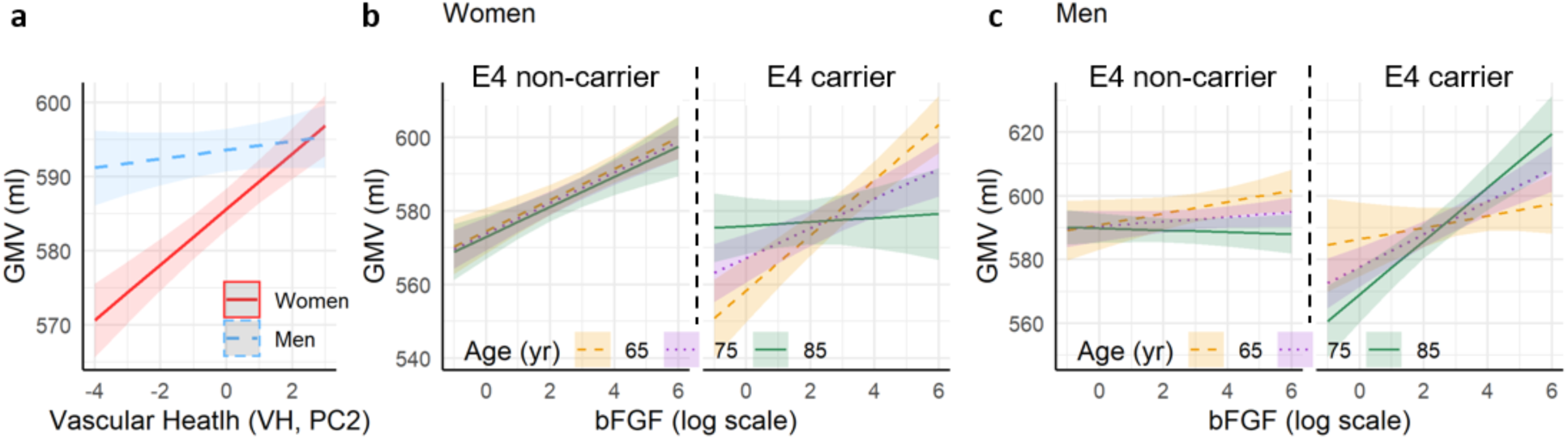
Association of Vascular Health with grey matter volume. The LMM analysis that models the association of Vascular Health (PC2) with GMV show a significant interaction between VH and sex to explain GMV (p = 0.03) (**a**). The LMM model with bFGF as predictor shows a significant interaction between bFGF, age, sex and APOE4 genotype to explain GMV (p = 0.023). The prediction of GMV for three ages (65, 75 and 85 years), and E4 non-carrier and carrier Women and Men are shown in **b** and **c** respectively. Shadow ribbon shows the standard error.

**Table 3.**
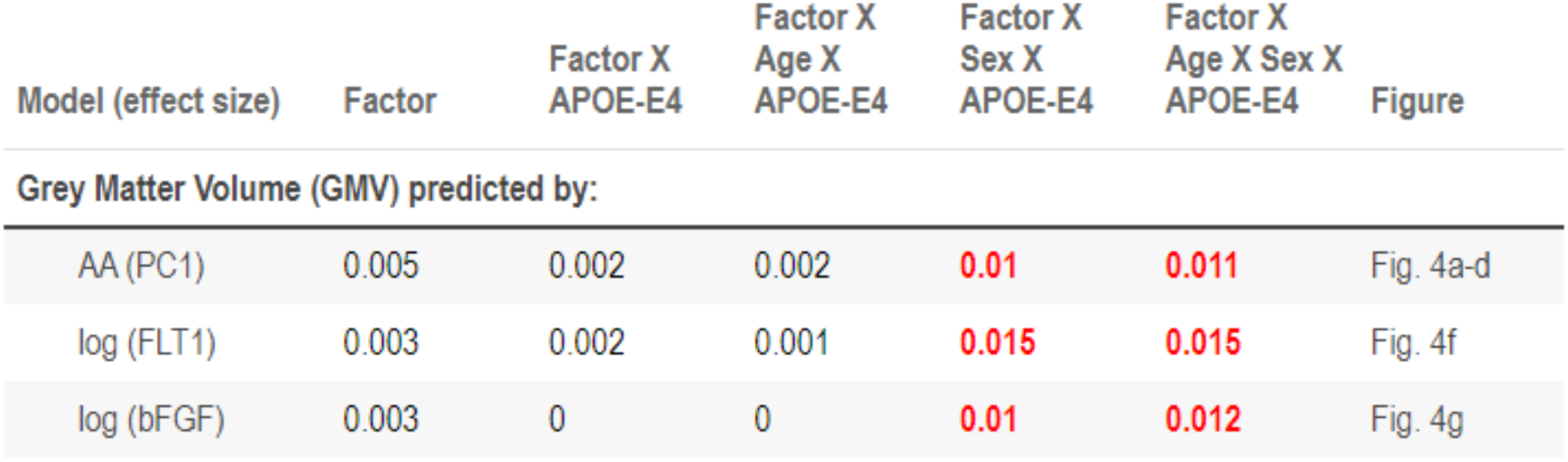
Summary statistics of interpreted models with APOE associations. Effect size (partial eta square) is shown for each association, with significant (p < 0.05) effects marked in red. “-“ indicates that the specific term as not tested. X indicates interaction. Factor makes reference to the specific predictor denoted in the first column

We find that higher values of VH are strongly associated with higher values of GMV in women, but to lesser extent in men (**Fig. 5a**). Congruent with the emerging role of bFGF and associated VH as potential markers of healthy brain aging or resilience to pathology, we find a positive association between bFGF and GMV in both men and women (**Fig. 5b-c**).

Ultimately, pathological aging results in cognitive impairment(*22*). Thus we investigated the association of plasma markers with executive function (**Fig. 6**), a well suited measure for VCID(*6*). We find that AA interacts with sex and age in predicting executive function (**Fig. 6a**). In line with results discussed above, we again find that higher levels of AA are associated with better executive function in younger women and lower executive function in older women, with an inflection point at age 78. Men of all ages show an inverse association between AA levels and executive function. We also find that PlGF is inversely associated with executive function, with higher levels in plasma associated with lower executive function, while FLT1 shows a similar pattern to AA with higher levels associated with lower executive function in older woman but not men (**Fig. 7**), and shifting in directionality around age 75. For VH, we find that higher levels are associated with higher executive function dependent on age, regardless of sex.

**Figure 6.**
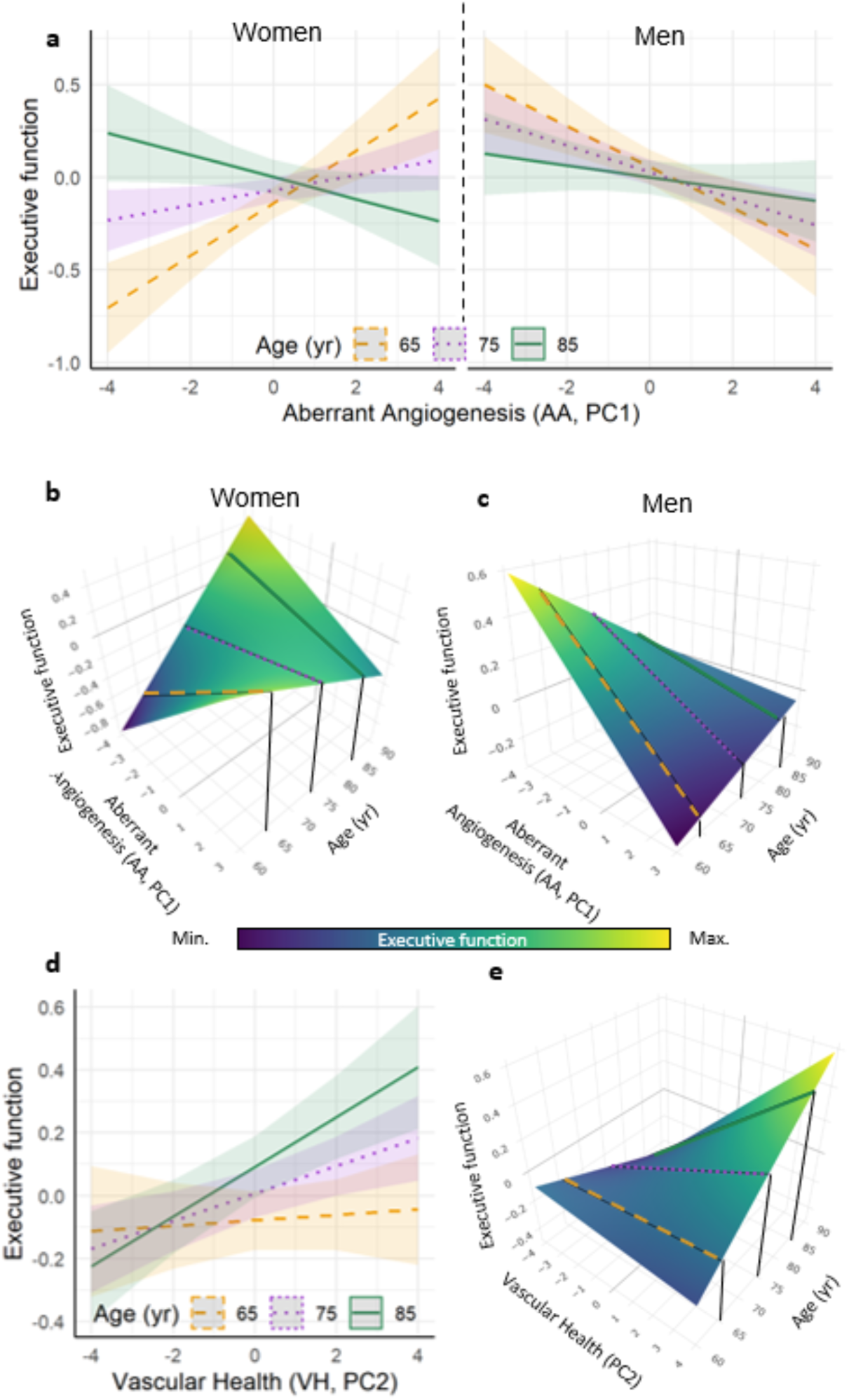
Association of executive function with Aberrant Angiogenesis and vascular health. The LMM model for the prediction of the executive function by AA results in a significant interaction between AA, age and sex (p = 0.021) (**a**, **b**, **c**). The prediction of executive function for three ages (65, 75 and 85 years) is shown in **a**, while the surface plot for the continuous interaction of age and AA is shown in **b** for Women and **c** for Men. In the case of the LMM model with VH as a predictor, there is a significant interaction between VH and age (p = 0.037) as shown in **d** and **e.** The interactive plot for **b**, **c** and **e** can be found on the supplementary material (supplementary interactive figures). The graphs show the predicted value (bold line or surface) and standard error (shadow ribbon).

**Figure 7.**
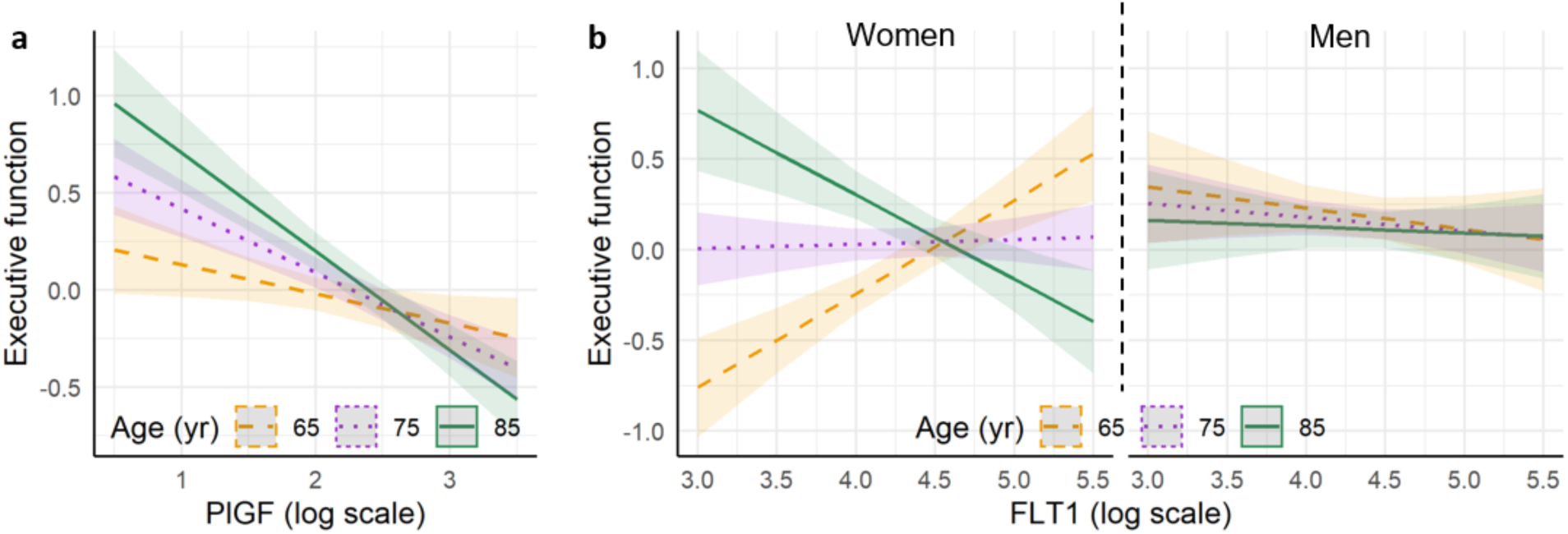
Association of executive function with PlGF and FLT1. The LMM model for the prediction of the executive function by PlGF results in a significant interaction between PlGF levels and age (p = 0.048) (**a**). In the case of the LMM model with FLT1 as a predictor, there is a significant interaction between FLT1, age and sex (p = 0.016) as shown in **b.** The graphs show the predicted value (bold line) and standard error (shadow ribbon). The prediction of executive function for three ages (65, 75 and 85 years) is shown.

#### Amyloid-independent contribution of plasma angiogenesis markers to cognition

We next tested whether angiogenesis markers have an amyloid-independent association with mild cognitive impairment, as a first step toward deciphering whether the associations are relevant to brain dysfunction and degeneration in aging, independent or in addition to AD-associated neurodegenerative disease. We performed a simple cross-sectional approach and compared levels of angiogenic predictors among amyloid positive and negative groups with and without cognitive impairment. Here, amyloid positivity is used as a marker of AD disease state.

W hypothesized an independent association of angiogenesis factors with cognitive impairment, with levels of AA, PlGF and FLT1 expected to be highest in the amyloid negative mild cognitive impairment (MCI) group. In these cross-sectional analyses, we used Amyloid PET status and CDR to classify disease state and stage, respectively (**Fig. 8**). For models looking at derived principal components, we find that in cognitively normal individuals, the levels of AA did not significantly differ in amyloid positive versus negative states. However, the amyloid negative MCI group (MCI-) demonstrate significantly higher levels of AA in comparison to amyloid positive MCI group (MCI+), rejecting our null hypothesis. The VH model was not significant. With respect to individual markers, FLT1 results echoed those obtained with AA, but the PlGF model was not significant. Interestingly, bFGF levels were found to be significantly higher in CN compared to MCI groups, regardless of amyloid status, in full alignment with results reported above, and suggestive once again that higher plasma bFGF levels may reflect a better brain health state.

**Figure 8.**
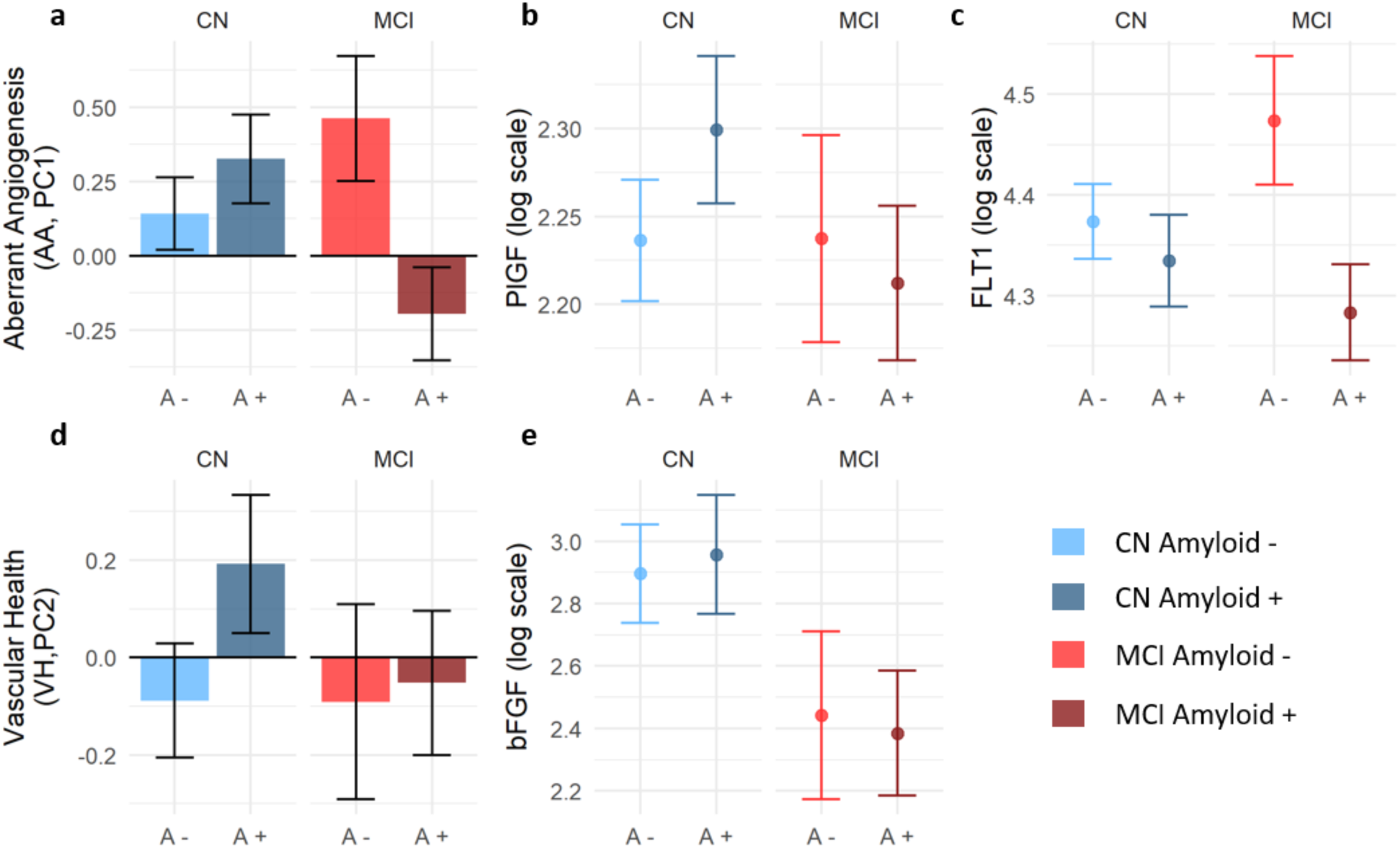
AA is elevated in cognitively impaired individuals with negative amyloid PET scan. In this cross-sectional analysis groups were formed using amyloid PET and CDR into account for disease state and clinical stage respectively. Linear models adjusted by age and sex followed by ANOVA were used to analyze the significance of amyloid status, the clinical stage and their interaction of the levels of AA, VH and the different markers. A significant interaction was observed for AA (**a**; p = 0.005). No significant terms were observed for PlGF (**b**), while a significant effect of amyloid was observed for FLT1 (**c**; p = 0.048). In the case of VH (**d**) no significant effects were detected, while bFGF was significantly different between clinical stages (**e**; p = 0.007) irrespective of amyloid status. In red amyloid negative; CN, cognitively normal; MCI, mild cognitive impairment.

Overall, these grouped, cross-sectional analyses suggest that dysfunctions in angiogenesis may mark independent pathological contribution(s) to cognitive impairment in aging and that plasma markers of angiogenesis may be promising biomarkers to investigate in VCID, independent, additive, or perhaps even synergistic with classical AD-associated pathologies.

## DISCUSSION

Well-orchestrated angiogenesis is fundamental to organismal health(*1*). Therefore, age-related aberrant angiogenesis may be a feature of numerous age-associated diseases(*23, 24*). To date, cross-sectional studies have reported increased levels of some angiogenesis factors in neurodegenerative diseases(*25–27*). In this paper, we map the longitudinal trajectory of angiogenesis factors in relation to brain aging trajectories employing a non-invasive molecular approach from plasma samples. We made several discoveries. At a fundamental level, we demonstrate strong associations between levels of plasma angiogenesis factors and brain health outcomes, suggesting that angiogenesis is highly relevant to diverse brain aging trajectories. Importantly, we uncover sex dimorphic and age-dependent directionality in these associations, with inflection points and change in directionality around age 75, suggesting that sex and age are important biological variables to consider in studies of angiogenesis. Sexual dimorphism has been well described in aging(*28–30*) and neurodegenerative disease literature(*31, 32*), however this is the first report of sex effect on the association of angiogenesis factors with brain aging outcomes in humans.

We extracted PCs in addition to the investigation of select markers of interest. We found that AA (Aberrant Angiogenesis, PC1), FLT1, and PlGF demonstrate age-associated increases, while VH (Vascular Health, PC2) and bFGF demonstrate age-associated decline in levels in both sexes. These were the first indications that higher AA, PlGF, and FLT1 levels may reflect pathological states in brain aging, and that higher levels of VH and bFGF could be reflecting healthier brains states. Next, we investigated associations of these markers with GMV and executive function, as two classical measures of structural and functional brain aging. Higher scores on executive function and higher GMV reflect better brain health in aging(*29*). Indeed, we found higher levels of AA and FLT1 to be associated with lower executive function in all groups except younger women, where higher and increasing levels were associated with higher executive function. A similar pattern was noted for AA, FLT1, and PlGF with GMV, with the addition of older men also demonstrating a surprising direction of association, akin to younger women. These sex dimorphic findings suggest future directions for this work. There may be a role for sex hormones in the biology underpinning the observed associations. It is also possible that, akin to proinflammatory cytokines that can have anti-inflammatory effects depending on the cells they are acting on and other molecular factors within the cellular microenvironment, the angiogenesis factors we quantified may have beneficial effects, be it on angiogenesis or other cellular processes benefiting cognition, based on the sex and age-dependent physiological states. It should be noted that, since our study is the first of its kind, absence of significance in any tested associations cannot be interpreted as lack of biological link. We used the largest cohort available for the study but may be underpowered for associations with smaller effect sizes. To understand the mechanistic meaning of these associations and identify therapeutic opportunities with respect to prevention or treatment of cognitive impairment, larger scale, multicenter follow-up studies are needed to get at causality.

We discuss several possible interpretations for these findings. First, Grunewald and colleagues recently demonstrated that FLT1 undergoes an age-related shift in alternative splicing, with increase in a soluble form (sFLT1), which can bind and sequester VEGFA, functioning essentially as a decoy(*1*). In absence of compensatory increases in VEGFA, sequestration by sFLT1 can interfere with and impair physiological angiogenesis and vascular plasticity. In our current cohorts, we did not note an age-associated increase in VEGFA, that could putatively counteract increasing levels of sFLT1. The antibodies used for quantification of FLT1 in this study would recognize both bound and secreted forms. Although the membrane bound FLT1 could become detectable in blood with cell turnover, it is unlikely to be the major source of signal in absence of acute injury. Therefore, we reason that age-associated increase in levels of FLT1 may be due to this age-related shift from bound to secreted forms. It is also interesting that among single factors, FLT1 demonstrated a clear inverse association with executive function in older women, and a positive association in younger women. It is possible that too much sFLT1 becomes problematic in old age, but that this isoform has a physiological function in fine-tuning angiogenesis in younger women, since uncontrolled angiogenesis can have pathological consequences, such as rise of dysfunctional, leaky blood vessels(*14, 33–35*). Second, the most potent version of VEGFA ligand is as a homodimer, signaling through membrane bound FLT1(*36, 37*). The ligand, PlGF can form heterodimers with VEGFA, leading to an attenuated activation of angiogenesis in comparison to VEGFA homodimers, with additional consequences of a pathological angiogenic signaling(*37–39*). These putative biological mechanisms could explain the inverse associations noted between levels of PlGF with executive function, regardless of age and sex. In fact, prior studies have demonstrated pathological effects of PlGF and identified this factor as a potential target for inhibition of pathological angiogenesis without interfering with the necessary, required physiological angiogenesis, so important for organismal health(*40*). However, how PlGF is regulated in the aging brain, and whether it has both physiological and pathological effects, remains to be determined on a mechanistic level.

We take these results to indicate that higher levels of AA, PlGF and FLT1 are indicative of lower brain health state in aging, akin to pathological angiogenesis in retinal disease and cancer(*36, 41*), with a change in directionality of associations noted in mid 70s and with sex effects that need to be considered. With respect to health-associated angiogenesis markers, prior work has shown that bFGF can induce angiogenesis through the promotion of endothelial cell growth and enhanced cell division. In addition, as a neurotrophic factor, bFGF can contribute to neuronal resilience in the setting of vascular injury, neuroinflammation, and oxidative stress. However, whether systemic elevation in these levels have health benefits in the aging human brain or levels are highly correlated with other molecular processes of relevance to brain health remains to be determined on a mechanistic level.

In an aging population without known neuroimmunological disorders, such as multiple sclerosis, or acute vascular injury (stroke), higher levels of WMH are typically interpretated as a sign of underlying chronic small vessel disease, with dysfunction of the blood-brain barrier and reactive gliosis(*42*). In our study population, we noted a significant association between levels of AA and WMH, with a significant sex effect. We also noted a significant association between WMH and bFGF, with changing directionality across age. Studies have shown that bFGF can attenuate astrocyte activation by reducing the expression of glial fibrillary acidic protein (GFAP)(*43*). Moreover, in *in vitro* studies, expression of bFGF counteracts the many deleterious effects that amyloid beta exposure can have on endothelial cells and the function of angiogenesis. It remains to be determined whether increasing levels of bFGF can benefit brain health measures. If bFGF attenuates astrocytic activation, it could have direct and indirect mechanisms of action, potentially mediated through astrocytic phenotypes(*44*).

Overall, impaired, reactive, or dysregulated angiogenesis may be an early molecular pathology contributing to adverse cognitive aging, or an amplifier, accelerator of brain dysfunction in aging, triggered by age-associated biochemical and functional changes. Relation and interactions with other age-associated brain pathologies, such as decreased brain clearance and accumulation of amyloid beta (Aβ) species, with impact on endothelial cells, has been studied *in vitro* and should be investigated *in vivo*. In *in vitro* and *in vivo* model systems, it has been shown that all amyloid beta (Aβ) species can inhibit angiogenesis (Merkulova-Rainon et al., 2018; Paris et al., 2004; Solito et al., 2009, Parodi-Rullan 2020). However, in the living human brains affected by aging and Alzheimer’s disease, it remains to be determined whether angiogenesis is inhibited, leading to decreased perfusion, capillary density, and inadequate response to neuronal demand, or whether it is dysregulated and uncontrolled, leading to formation of structurally and functionally abnormal leaky blood vessels that don’t appropriately incorporate into the neurovascular unit, or both. Ultimately angiogenesis may be dysregulated in a disease state and stage specific manner with levels of angiogenesis factors representing diverse molecular processes and dysfunctions of vascular cells and non-vascular cells. Our study provides compelling evidence in support of the utility of plasma levels of angiogenesis factors in studies of molecular drivers of cognitive aging. Future studies should dissect mechanisms and ideal interventions on the multitude of pathways captured by these markers.

In this work, we provide evidence to suggest that interventions to boost or modulate angiogenesis may benefit brain aging, similar to effects shown on other organs(*1*). Similarities and differences between brain and systemic vascular physiology and pathology is under investigation, however, circulating angiogenesis factors could plausibly reflect a general state of vascular health or dysfunction. By reaching endothelial cells throughout the organism, angiogenesis factors could act on numerous vascular beds, including the brain. With vascular pathologies contributing to brain aging and neurodegenerative disease, the modeling approach we adopted in this study provides a blueprint for investigation of plasma biomarkers of vascular neurodegenerative disease, with great promise for future diagnostics and clinical trials interventions.

## METHODS

### Study Participants

Participants were enrolled into the MarkVCID study, selected from ongoing longitudinal studies of brain aging and dementia at two NIH funded Alzheimer’s Disease Research Centers (ADRC), namely the Memory and Aging Center at University of California San Francisco and University of California Davis. Assessments were similar across both sites, with informant interview for assessment of functional status (Clinical Dementia Rating, CDR), neuropsychological testing, and comprehensive physical and neurological examination, questionnaires and history. Biospecimen had been collected at each site, processed similarly, according to ADRC protocols, aliquoted and stored at −80°C until used. All study participants provided informed consent and the study protocols were approved by the UCSF and UCD Human Research Protection Program and Institutional Review Board. Research was performed in accordance with the Code of Ethics of the World Medical Association.

### 2.2. Cognitive Testing and Functional Assessments

The Clinical Dementia Rating scale (CDR) is a global assessment of functional status and clinical disease severity with six subdomains obtained via study partner interviews(*45*). For the derivation of this score, six subdomains are scored on a five-point scale (0/0.5/1/2/3), with higher scores indicating greater disease severity. A CDR sum of the boxes (subdomains) is calculated (CDRsob), which sums the subdomain scores (range: 0-24). From the CDRsob, a global score (CDR) is calculated that is a weighted summary of the six subdomain scores.

Participants completed several neuropsychological tests of executive functions, including a modified trail-making task (Kramer et al., 2003), Design Fluency (Filled Dots Condition) from the Delis–Kaplan Executive Function System (D. Delis et al., 2001), lexical fluency (Birn et al., 2010), Stroop color–word interference (Stroop, 1935), and a backward number span task (Kramer 2003; weintraub). Trail making test scores were logarithmically transformed to normalize the data. Performances on neuropsychological tests of executive function were summarized into a single composite score that was calculated as described in detail elsewhere(*46*). Briefly, scores on each test were converted to z-scores relative to the mean and standard deviation of a control sample recruited at UCSF. The executive composite was created by averaging the available z-scores for at each observation. The composite was created for participants as long as they completed three of the five measures.

### Neuroimaging Evaluation

#### MRI Acquisition

Subjects were scanned on a Siemens Prisma 3T and Trio scanners at the UCSF Neuroscience Imaging Center and University of California Davis, respectively. A T1-weighted Magnetization-prepared rapid gradient echo (MP-RAGE) structural scan was acquired in a sagittal orientation, a field-of-view of 256 x 240 x 160 mm with an isotropic voxel resolution of 1 mm3, TR = 2300 ms, TE = 2.9 ms, TI = 900 ms, flip angle = 9°. The T2 fluid attenuated inversion recovery (FLAIR) acquired in the axial orientation, field-of-view = 176 x 256 x 256 mm, resolution 1.00 x 0.98 x 0.98 mm3, TR = 5000 ms, TE = 397 ms, TI = 1800 ms.

#### MRI Processing and Analyses

De-identified digital information was transferred from UCSF to UCD using secure and HIPAA complaint DICOM server technology. UCSF and UCD images were processed by the Imaging of Dementia and Aging (IDeA) lab at UC Davis and full imaging protocol details are reported in prior publications(*47–50*). In brief, WMH quantification was performed on a combination of FLAIR and 3D T1 images using a modified Bayesian probability structure based on a previously published method of histogram fitting. Prior probability maps for WMH were created from more than 700 individuals with semi-automatic detection of WMH followed by manual editing. Likelihood estimates of the native image were calculated through histogram segmentation and thresholding. All segmentation was initially performed in standard space resulting in probability likelihood values of WMH at each voxel in the white matter. These probabilities were then thresholded at 3.5 SD above the mean to create a binary WMH mask. Further segmentation was based on a modified Bayesian approach that combines image likelihood estimates, spatial priors, and tissue class constraints. The segmented WMH masks were then back-transformed on to native space for tissue volume calculation. Volumes were log-transformed to normalize population variance.

### Plasma marker measurement

Plasma samples used for this study had not undergone any freeze-thaw cycles prior to analysis. All analyses were conducted by a board-certified laboratory technician blinded to study design. All assays were run using the V-Plex plates from Meso Scale Discovery (MSD), a high performance electrochemiluminescence (HPE) technology. The multiplex arrays were analyzed with a MESO QuickPlex SQ 120 imager (MSD, Rockville, MD) and Discovery Workbench v4.0 software. Concentrations were obtained in duplicate per each sample in accordance with the manufacturer’s protocol and coefficients of variance (CV) calculated. Samples with CV >10% were censored from analyses.

### Unsupervised learning

We performed unsupervised pattern detection using a combination of missing data analysis, data imputation and non-linear principal component analysis (NL-PCA). This workflow was performed twice, one for extracting patterns among angiogenesis markers and one for extracting the associations among vascular factor and disease variables. Data was extracted for each visit with available data for each patient.

*Missing data analysis and imputation:* both the angiogenesis markers and the vascular factor dataset contained missing values. For some visits some of the angiogenesis markers were out of range or the CV was higher than 20%. In the case of the vascular related disease variables, in some patients the data were only available for some of the visits. This created missingness due to the fact that medical history data collection might not happen at every visit. Given that the vascular disease variables are associated to past history, we performed last observation carry forward imputation (LOCF). Missing pattern visualization were obtained using the naniar (Tierney et al., 2020) R packages (**supplementary fig. 1**). Test for whether the missing data was Missing Completely At Random (MCAR) was performed using the TestMCARNormality() function from the MissMech package(*51*). The results of the test indicated that the MCAR hypothesis is rejected (p < 0.001). Multiple imputation was performed using predicting mean matching method available in the mice(*52*) R package, setting the number of imputations to m = 20. A list of 20 complete datasets were then obtained and aggregated using the median value for each observation to run NL-PCA.

*Unsupervised pattern detection with NL-PCA:* We performed robust NL-PCA by optimal scaling and alternated least squares(*53*). We used the implementation of the *princals* algorithm in the Gifi R package, setting the variables to be ordinally restricted to allow only for monotonic transformations in b-spline link functions of degree 2 and 3 internal knots at the tertials data points. The number of components to keep was determined by the permutation-based test of the Variance Accounted For (VAF) with 500 permutations, implemented in the R syndRomics package(*54*). The first two components were statistically considered not random, indication of meaningful patterns. The third component was also statistically significant from randomness, however stability analysis (see below) suggested an instable pattern and therefore PC3 was not selected. In the case of the vascular disease related variables, the first 3 PCs were selected as indicated by the permutation test and stability analysis. We used these components for further interpretation and analysis. Visualization of the NL-PCA results was performed using the R syndRomics package.

*Component stability analysis:* to test whether the components or patterns determined by NL-PCA are stable under resampling conditions, a way to estimate the robustness and reproducibility of the solution, we used bootstrapping as implemented in the R syndRomics package. We used 500 bootstrapped samples and derived several metrics of component similarity (Pearson correlation, coefficient of congruence, root mean standard error and Cattell’s s-index), as well as the distribution of component loadings. From these, the average and the 95% confidence interval across the 500 resamples were obtained. Our results indicate high stability of the results for angiogenesis PC1 and PC2, as well as vascular PC1, PC2 and PC3, suggesting that the patterns are reproducible and generalizable (**supplementary fig. 1-2**). In addition, we studied the sensitivity of the NL-PCA solution to missing data and multiple imputation by comparing the NL-PCA solution from the 20 imputed datasets to the aggregated NL-PCA. Component similarity metrics indicated that PC1 and PC2 were stable under the multiple imputation model (coefficient of congruence of 0.996 and 0.991, respectively), while PC3 showed more variations across multiple imputed datasets (coefficient of congruence of 0.91).

### Regression analysis with linear mixed models

In order to study the association of the derived principal components and individual markers to key variables, we performed regression analysis. Due to the repeat assessments within a participant across different visits over time, linear mixed models (LMM) where fitted to deal with the intra-subject correlation structure by setting participant as random intercept term. All other predictors were specified as fixed effect term. LMM were fitted using the lmer() function in the lme4 R package(*55*). An F-test with a type III sum of squares was used for testing the hypothesis of whether a given fixed term had a statistically significant contribution in explaining the variance in the response variable for the LMM fitted model. For studying the association of the PC scores and markers with demographics variables, a LMM model was fitted with the PC score or marker as response variable and age, sex and their interaction as fixed effect terms. For the study of whether each PC or marker is a predictor for the imaging, cognitive of vascular variables, a different LMM model was fitted, adjusting by age and sex. To understand the potential moderation effect of age and sex on each marker and PC association with the response, we first fitted a triple interaction model between the predictor of interest, age and sex. If the triple interaction was not significant, with a small F value and high p value, the model was simplified to include only double interactions, and simplified further to an additive-only model (no interaction terms) in case no interaction was significant. To study the effect of APOE4 genotype on the association of the PCs or markers with the response variable, we included APOE4 status as a new term in the model, including all possible interactions. Model simplification was carried as described above. The final model to interpret was the most parsimonious model that could not be simplified further. An alpha of 5% (p = 0.05) was considered a cut off for statistical significance. The results of all fitted models can be found in the supplementary tables.

We investigated the relationship of the predictors further by using interaction plots of the predicted value from the LMM fitted models. In the case of bivariate continuous interaction between the main predictor of interest (PC score or marker) and age, three predicted regression lines of the response variable were obtained for the range of the main predictor at three different ages (65 years, 75 year and 85 years) which corresponds to the rounded mean and ± standard deviation of age. Prediction interaction plots with the regression line and its ± standard deviation band were obtained using the sjPlot or the emmeans R packages. Since continuous bivariate interactions produce a prediction surface, we generated 3D prediction surface plots when needed using a custom script and the ggplot2 and plotly R packages. A snapshot of these is included in the figures. The interactive 3D plots can be explored in the supplementary figures.

### Cross-sectional analysis

For the cross-sectional analysis of amyloid status and clinical stage associations we selected the first visit with available marker data for each patient. Amyloid status (positive or negative) was determined using standardized PET cut off values. Clinical stage was determined by CDR to classify disease state into functional impaired (MCI) or unimpaired (CN). Thus, we defined four categories as follows: CN-, CN+, MCI-, MCI+. Linear models with PC scores or the marker of interest as the response variable and amyloid status, clinical stage and the interaction between them where fitted, adjusting for age and sex. An F-test was used for testing the hypothesis of whether a given term had a statistically significant contribution in explaining the variance in the response variable for the fitted model. An alpha of 5% (p = 0.05) was considered a cut off for statistical significance. The results of all fitted models can be found in the supplementary tables.

## Acknowledgements

This work was funded by the National Institute on Aging and Department of Veterans Affairs IK2CX002180, Larry L. Hillblom Foundation 2019A012SUP and New Vision Research (to F.M.E.). Participant recruitment and data collection at UCSF was funded by the MarkVCID (UH3NS100608) and Hillblom Aging Network for the Prevention of Age-Associated Cognitive Decline study grants (2140-A-004-NET) (to J.K.) and NIH-NIA ADRC (P50AG023501) and PPG (P01AG019724) (to B.L.M.). Participant recruitment and data collection at UCD was funded by P30AG010129 and P30AG072972 (to C.D.)

## Author contributions

All authors provided critical input, reviewed, and edited the manuscript. Conceptualization: FME. Research participant selection: FME, JHK, CD, and KBC. Data collection: FME, KBC, and LG. Analytical plan: FME, ATE, ARF. Figures and Tables generated by ATE. Cognitive test composites were created by AMS. PM and CD processed all brain imaging. FME drafted the manuscript with critical input from ATE and edits from all co-authors. JHK, CD, and BML led the creation of cohorts used in the study.

