## Supplementary figures and images for "Sexually dimorphic differences in angiogenesis markers predict brain aging trajectories"

### Supplemental figures

#### Supplementary Figure 1

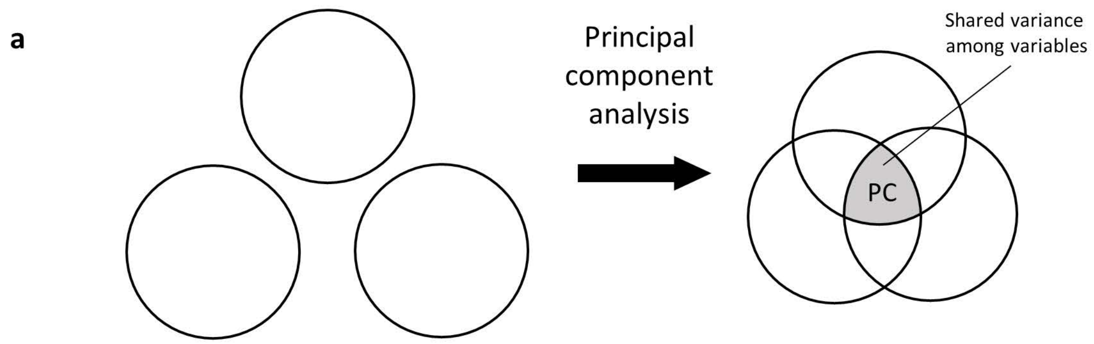


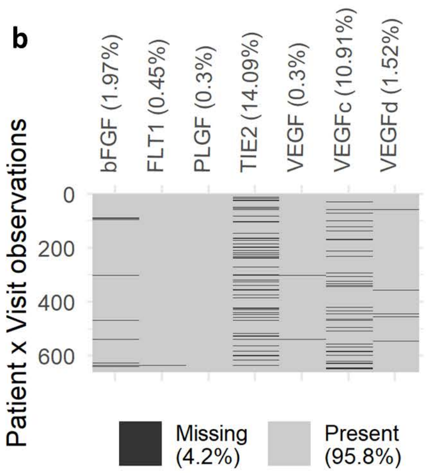


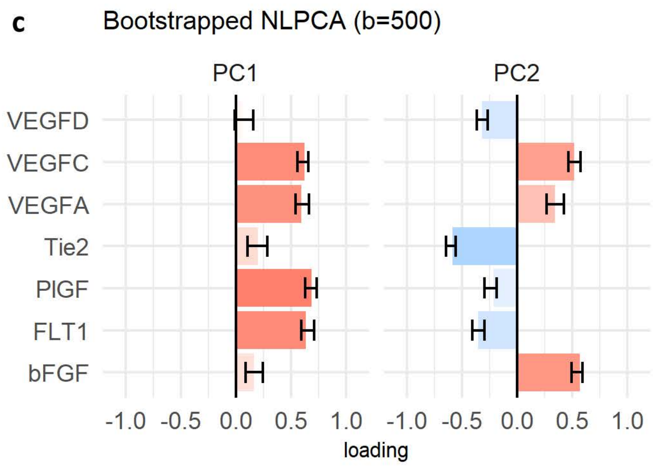
